# Structures of LIG1 uncover the mechanism of sugar discrimination against a ribonucleotide at 3’- and 5’-end of the nick DNA

**DOI:** 10.1101/2022.08.26.505476

**Authors:** Qun Tang, Mitchell Gulkis, Melike Çağlayan

## Abstract

Human DNA ligase I (LIG1) is the main replicative ligase that seals Okazaki fragments and finalizes DNA repair pathways by joining canonical 3’-OH and 5’-P ends of the nick DNA in a three-step ligation reaction. Ribonucleotides can be misincorporated by DNA polymerases resulting in a nick with 3’-ribonucleotide while RNase H2 mediated cleavage leaves a nick harboring 5’-ribonucleotide during ribonucleotide excision repair. However, how LIG1 surveils DNA ends with a “wrong” sugar at atomic resolution is unknown. Here, we determine X-ray structures of LIG1/nick DNA complexes with 3’- or 5’-single ribonucleotide during different stages of the ligation reaction. Our LIG1/5’-rG:C structure reveals a global conformational change, which discriminates against 5’-RNA/DNA junctions at the initial step when the ligase-AMP intermediate is formed. Furthermore, we capture LIG1/3’-RNA-DNA heteroduplexes that are tolerated at the active site where AMP is transferred to nick DNA (step 2) and final phosphodiester bond formation occurs (step 3). Finally, we demonstrate the mutagenic and defective ligation of the nick DNA with 3’- and 5’-ribonucleotide, respectively, *in vitro*. Together, these results uncover how LIG1 encounters ribonucleotides embedded into genome during nuclear replication and the last step of DNA repair pathways to maintain genome integrity.

---

Mammalian DNA ligases contain a conserved three domain core architecture consisting of the oligonucleotide binding (OBD), the adenylation (AdD), and the DNA binding (DBD) domains, which seal 3’-hydoxyl (OH) and 5’-phosphate (P) termini of the nick DNA in a conserved ligation reaction involving the three consecutive chemical steps^1,2^ (Extended Data Scheme 1): the formation of the DNA ligase-adenylate intermediate (LIG-AMP) in step 1, subsequent transfer of AMP moiety to the 5’-P end of the nick (DNA-AMP) in step 2, and final phosphodiester bond formation coupled to AMP release in step 3^3,4^. Despite these similarities, human DNA ligases exhibit different ligation fidelity which relies on the accuracy of DNA synthesis that requires a Watson-Crick base pair when DNA polymerase incorporates a correct nucleotide during DNA replication and repair^5,6^.

The concentration of ribonucleotide triphosphates (rNTPs) is much more abundant than those of deoxyribonucleotide triphosphates (dNTPs) within a cell, and therefore, their misincorporation into genomic DNA by DNA polymerases occur at higher frequencies than mismatch nucleotide insertions, making genomic ribonucleotides the most prevalent source of cellular DNA damage^7–10^. In human cells, ribonucleotide incorporation can occur during almost all cellular DNA transactions^11–17^, and discrimination against ribonucleotide incorporation by replicative and repair DNA polymerases is essential for the maintenance of genomic integrity^18–27^. Ribonucleotides embedded into genomic DNA can confound structural and chemical integrity of duplex DNA, increase its susceptibility to endogenous or exogenous damage, and lead to several types of genome instability such as replication blockage, mutagenesis, aberrant recombination, protein-DNA crosslinks, double-strand breaks, and chromosome alterations^28–31^.

Among the four known human DNA ligases, LIG1 plays a critical role in Okazaki fragment maturation during DNA replication and finalizes the last ligation step of most of DNA repair pathways^1–4^. In the first solved structure of LIG1, it has been shown in the ligation assays *in vitro* that the ligase displays a great discrimination against the DNA substrate when the 5’-phosphorylated strand is completely RNA, while it can ligate the RNA strand that is located at the upstream of the nick^32^. Yet, the mechanism of sugar discrimination by LIG1 at atomic resolution is entirely missing. Recently solved LIG1 structures demonstrated that the ligase employs a Mg^2+^-reinforced nick DNA-binding mode to ensure high fidelity ligation, and this accuracy is a critical determinant of faithful replication of the nuclear genome^33–35^. Furthermore, our structures revealed that LIG1 active site discriminates against mutagenic mismatches distinctly depending on the architecture of the 3’-terminus at the nick DNA^36^. The ligation efficiency for the ribonucleotides at positions near the ends of nick DNA has been also reported for DNA ligases from *Chlorella virus, Thermus thermophilus*, and *Saccharomyces cerevisiae*^37–41^.

Ribonucleotide Excision Repair (RER) is the primary mechanism through which ribonucleotides embedded into genomic DNA can be repaired^42,43^. The RER pathway is initiated by RNase H2-mediated incision of the DNA backbone at the 5’-end of the ribonucleotide, followed by strand displacement synthesis by pol δ, then flap removal by Flap Endonuclease 1 and/or Dna2, and finally nick sealing by LIG1^44–46^. Previous studies also demonstrated that LIG1 attempts to ligate the RNA-DNA lesion following RNase H2 cleavage, which results in the formation of abortive ligation products with a 5’-adenylated-ribonucleotide that can be resolved by Aprataxin (APTX)^47,48^. Despite of the fact that LIG1 involves in the RER process, how it surveils the RNA-DNA junctions harboring a 5’-ribonucleotide at atomic resolution remains unknown.

In the present study, we aimed to elucidate the mechanism by which human LIG1 discriminates against a single ribonucleotide at the 3’- and 5’-ends of nick DNA. Our LIG1 structures uncover a lack of proficient sugar discrimination and mutagenic ligation of nick DNA with a ribonucleotide at the 3’-end during different steps of ligation reaction (Extended Data Scheme 1). We also demonstrate a conformational change of LIG1 in the presence of 5’-RNA-DNA junction, which deters nick sealing during initial step. Overall, our findings provide a novel insight into the characterization of ribonucleotide selectivity by LIG1 on the downstream events of DNA replication and repair, demonstrating its ability to surveil ribonucleotide-containing ends of nick DNA distinctly depending on the position of an incorrect sugar.

## LIG1 fails to discriminate against a ribonucleotide at 3’-end of the nick DNA

We solved the structures of LIG1/nick DNA complexes containing rA:T, rC:G, and rG:C at the 3’-end (Fig. 1 and Extended Data Table 1) for LIG1^WT^ and LIG1^EE/AA^ that harbors E346A and E592A mutations resulting in the ablation of the high-fidelity site, which has been utilized in previous structures with non-canonical substrates^33–36^. The root mean square deviation (RMSD) of the LIG1 structures for 3’-rA:T, 3’-rC:G, and 3’-rG:C was no higher than 0.7 Å for all main chain atoms, which demonstrates that all LIG1 structures show similar global conformations (Extended Data Table 2). We observed the ligase active site in step 2 of the ligation reaction when the AMP moiety is transferred to the 5’-end of the nick DNA, forming a DNA-AMP intermediate, in the structures of LIG1^WT^/3’-rA:T and LIG1^EE/AA^ for 3’-rA:T, 3’-rC:G, and 3’-rG:C. Similarly, for the LIG1/nick DNA complexes containing canonical ends, LIG1^WT^/3’-dA:T and LIG1^EE/AA^/3’-dG:C structures showed step 2 (Fig. 1a-c,e,f,h). In addition, we captured LIG1^EE/AA^/3’-rA:T and LIG1^EE/AA^/3’-rG:C during the final step of the ligation reaction where the 3’-OH terminus attacks the 5’-P terminus downstream of the nick to form a phosphodiester bond (Fig. 1d,g). In the LIG1/nick DNA structures captured in step 2, the Fo-Fc maps at 3σ of the AMP moiety show a density for almost all the atoms in the AMP (Fig. 1a-c,e,f,h). In the step 3 structures, the Fo-Fc maps at 3σ of the AMP moiety are incomplete, suggesting that the AMP is released (Fig. 1d,g). The only difference in the crystallization conditions between step 2 (one day) and step 3 (three days) structures of LIG1 is the amount of time during which the ligase/nick DNA crystals were allowed to grow before freezing.

**Fig. 1.**
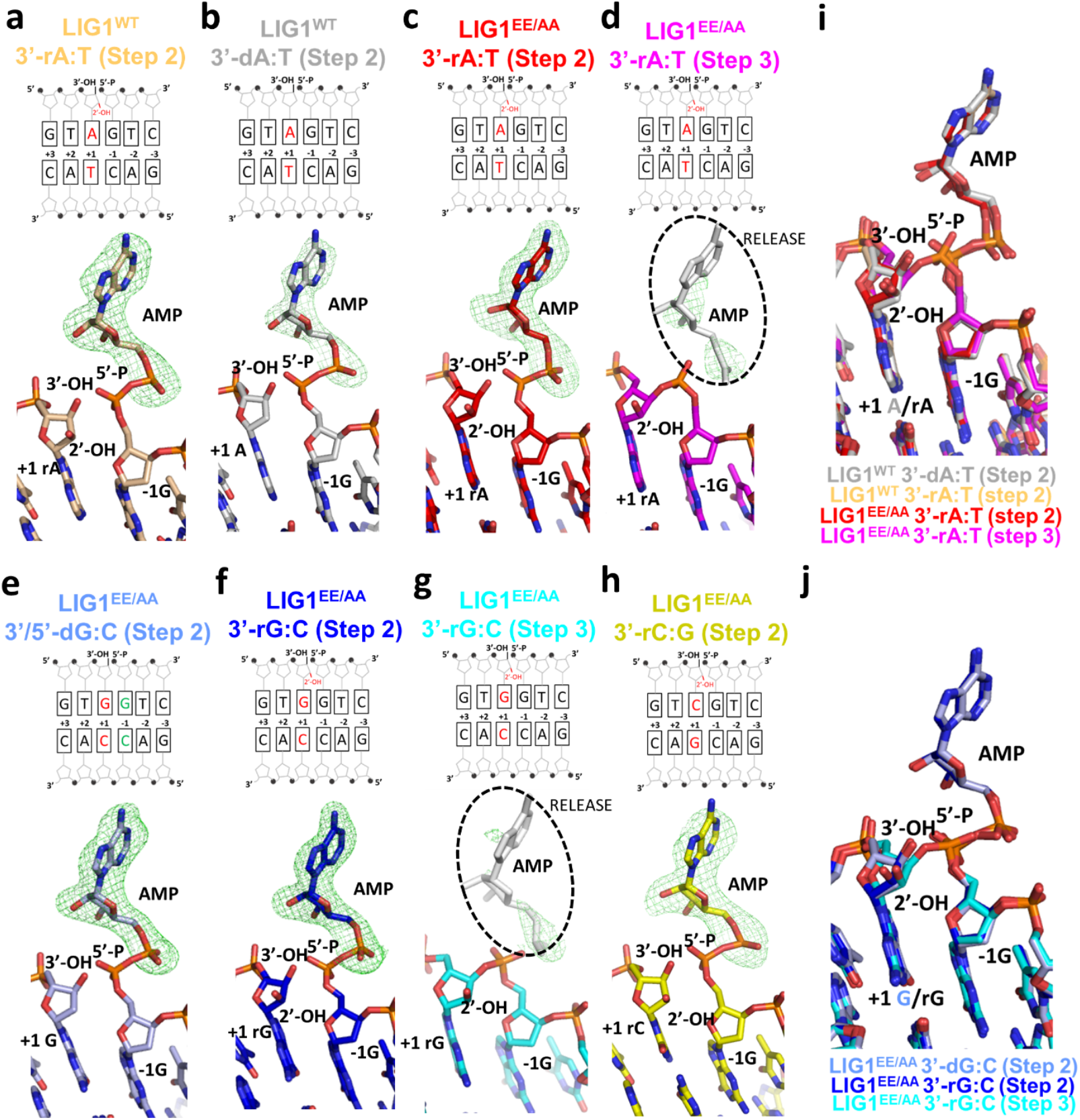
Structures of LIG1 bound to nick DNA duplexes containing 3’-ribonucleotide. Structures of LIG1^WT^ and LIG1^EE/AA^ in complex with nick DNA containing ribonucleotide or deoxyribonucleotide at the 3’-end are presented for LIG1^WT^/3’-rA:T (step 2, **a**), LIG1^WT^/3’-dA:T (step 2, **b**), LIG1^EE/AA^/3’-rA:T (step 2, **c**), LIG1^EE/AA^/3’-rA:T (step 3, **d**), LIG1^EE/AA^/3’-dG:C (step 2, **e**), LIG1^EE/AA^/3’-rG:C (step 2, **f**), LIG1^EE/AA^/3’-rG:C (step 3, **g**), LIG1^EE/AA^/3’-rC:G (step 2, **h**). Simulated annealing omit maps (Fo-Fc) of AMP are contoured at 3σ. Schematic view of the nick DNA shows the sequence of 3’- and 5’-terminus of the nick DNA substrate used in the LIG1 crystallization. In the step 2 structures, AMPs are built depending on the map, while in the step 3 structures, the AMPs do not exist because the maps of AMPs are incomplete. **i-j**, Differences in the positions of 3’- and 5’-terminus of the nick DNA are presented in the overlays of LIG1^WT^ and LIG1^EE/AA^ structures for the nick DNA duplexes containing 3’-dA:T and 3’-rA:T (i) and LIG1^EE/AA^ structures for the nick DNA duplexes containing 3’-dG:C and 3’-rG:C (j).

The superimposition of LIG1 structures showing step 2 (LIG1^WT^/3’-dA:T, LIG1^WT^/3’-rA:T, LIG1^EE/AA^/3’-rA:T) *versus* step 3 (LIG1^EE/AA^/3’-rA:T) of the ligation reaction demonstrates that the C3’ atom of the ribose on the 3’-end moves closer to the 5’-P at the nick in the ligase structures exhibiting a final phosphodiester bond formation (Fig. 1i). We observed a similar difference in the overlay of G:C structures showing step 2 (LIG1^EE/AA^/3’-dG:C, LIG1^EE/AA^/3’-rG:C) *versus* step 3 (LIG1^EE/AA^/3’-rG:C) (Fig. 1j). The distance between C3’ atom of the ribose and 5’-P was ~ 1.0 Å shorter (~ 3.5-4.0 Å and 2.6 Å which correspond to steps 2 and 3, respectively) in the step 3 structures of LIG1 (Extended Data Fig. 1). This is consistent with the formation of a covalent bond between the 3’- and 5’-ends of the nick. Furthermore, the comparison of the 2Fo-Fc maps at the 3’-OH and 5’-P ends of the nick demonstrated continuous and discontinuous maps in the LIG1 structures captured in the steps 2 and 3, respectively (Extended Data Fig. 1). Finally, we observed a Watson-Crick base pair at the 3’-end for all LIG1/nick DNA structures containing 3’-ribonucleotides and sugar pucker analyses demonstrate that the ribose adopts a C3’-endo conformation (Extended Data Figs. 2 and 3).

**Fig. 2.**
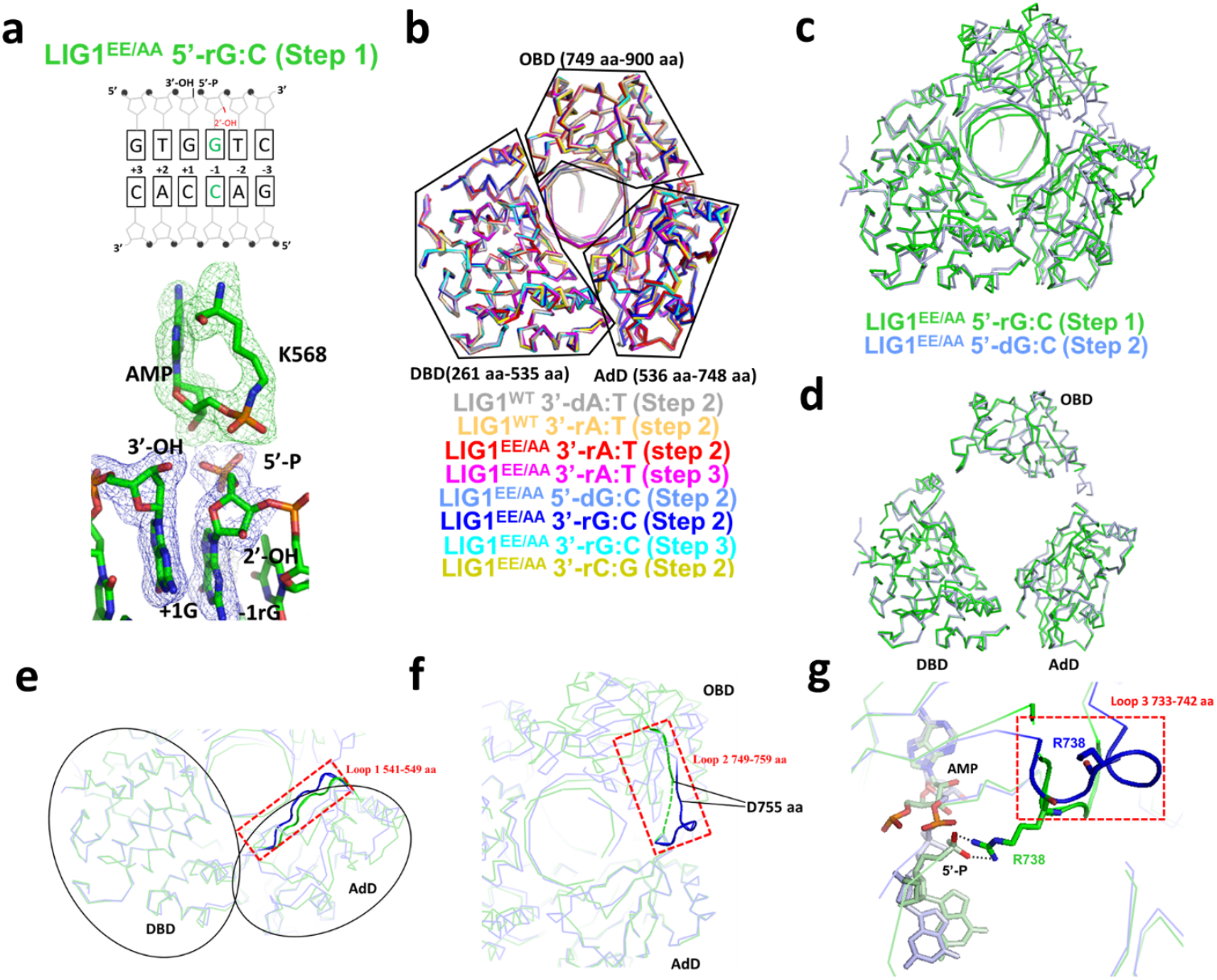
Structures of LIG1 bound to nick DNA duplexes containing 5’-ribonucleotide. **a**, Structure of LIG1^EE/AA^/5’-rG:C (LIG1^K568^-AMP intermediate, step 1). Schematic view of the nick DNA shows the sequence of 3’- and 5’-terminus of the nick DNA substrate. Simulated annealing omit map (Fo-Fc) of AMP is contoured at 3σ (green). 2Fo - Fc density map of 3’- and 5’-terminus of the nick DNAs are contoured at 1σ (blue). **b**, Overlay of LIG1^WT^ and LIG1^EE/AA^ structures for the nick DNA duplexes containing ribonucleotide or deoxyribonucleotide at the 3’-end are presented for the DBD, AdD and OBD domains of LIG1 encircling a nick DNA. **c-g**, Overlays of LIG1^EE/AA^ structures for 5’-rG:C and 5’-dG:C show overall (c) and individual ligase domains (d). The conformational changes in the loop 1 (red circle) residing in the AdD (e), and the loop 2 (red circle) residing in the OBD (f) domains. **g**, Conformational change at the loop 3 (red circle) shows a shift at the R738 side chain that interacts with 5’-P end of nick in the structure of LIG1^EE/AA^/5’-rG:C. The amino acid residue R738, AMP, and nick DNA are shown as a stick mode and all LIG1 are shown as a ribbon mode.

**Fig. 3.**
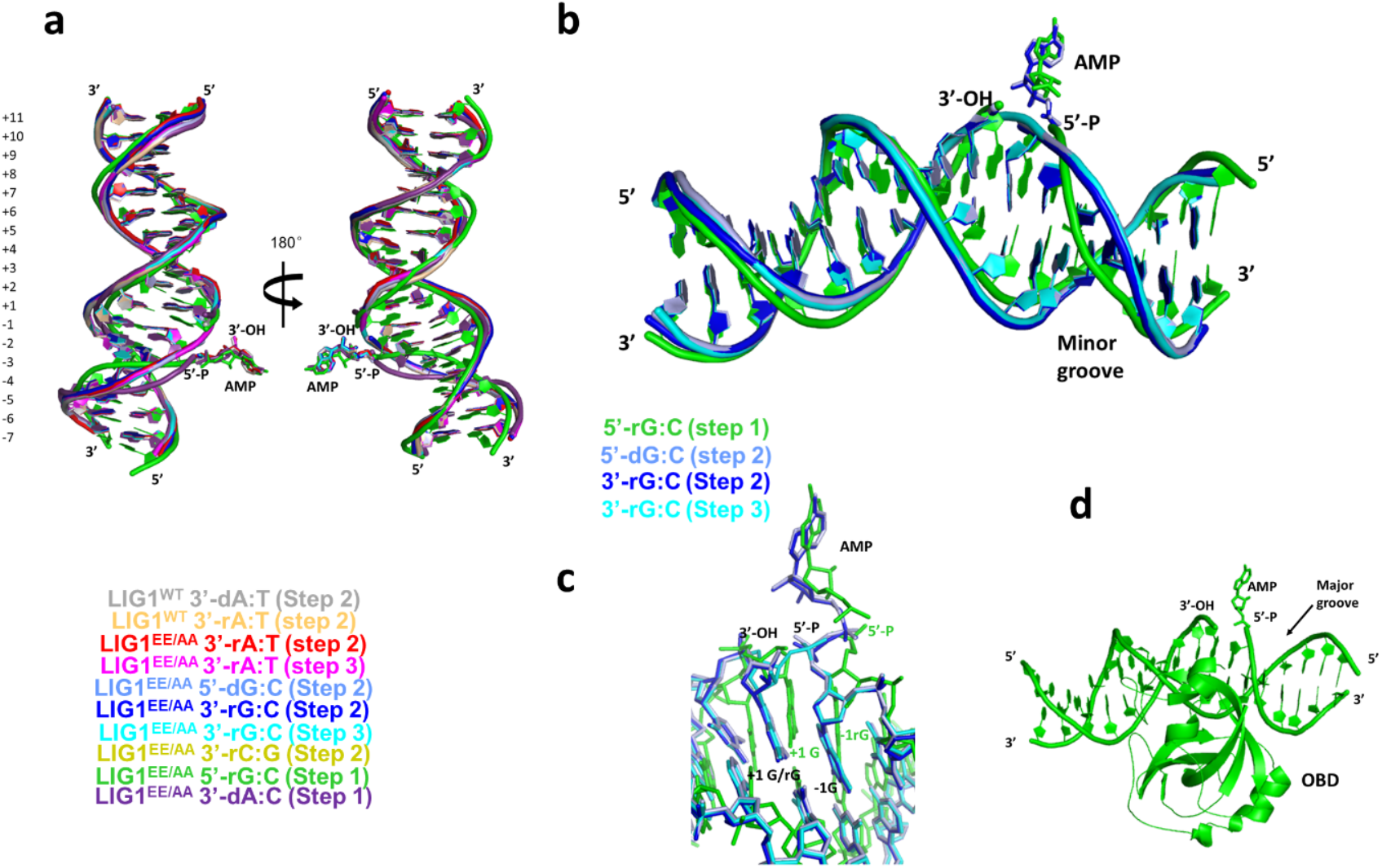
Structure of LIG1/nick DNA with a 5’-ribonucleotide uncovers a conformational change the minor groove. **a**, Superimposition of overall LIG1^WT^ and LIG1^EE/AA^ structures solved in this study shows a difference in the DNA conformation in LIG1^EE/AA^/5’-rG:C structure (green). **b**, Comparison of the nick DNA between LIG1^EE/AA^ structures captured in different steps of the ligation reaction; 5’rG:C (step 1), 3’-dA:C (step 1, 7SX5), 3’-dG:C (step 2), and 3’-rG:C (step 3); demonstrates a rotation of the 5’-P downstream of the nick DNA which widens the major groove downstream of the nick DNA and results in a narrowing of the minor groove in LIG1^EE/AA^/5’-rG:C structure. **c**, Overlay of LIG1^EE/AA^ structures for 3’-dG:C, 3’-rG:C, 3’-rG:C, and 5’-rG:C shows the conformation change at the 3’- and 5’-terminus of the nick DNA. **d**, Structure of LIG1^EE/AA^/5’-rG:C shows the conformation change in the OBD domain of the ligase interacting with nick DNA. OBD and nick DNA are shown as cartoon mode.

## LIG1 discriminates against RNA/DNA junctions harboring a ribonucleotide at the 5’-end

We captured the LIG1/nick DNA complex containing a 5’-ribonucleotide during the initial step 1 of the ligation reaction where an AMP moiety remains covalently bond to the active side chain of K568 (Fig. 2a). The superimposition of overall step 2 and 3 structures of the present study demonstrated that they are similar with the global conformation (Fig. 2b). When we overlayed the three domains (DBD, AdD, OBD) of LIG1^EE/AA^ for 5’-rG:C and 5’-dG:C structures individually, we found that the RMSD value was 1 Å, demonstrating no major conformational change within the individual domains (Fig. 2c,d and Extended Data Table 2). In previously solved structures of LIG1/nick DNA duplexes containing canonical, damaged (8-oxoG:A), and mismatched ends^32–36^, no global ligase conformational change has been reported and all the RMSD values have been found to be less than 1 Å. However, when we overlayed the entire catalytic core of LIG1^EE/AA^ for 5’-rG:C and 5’-dG:C, as well as our other LIG1 structures of 3’-ribonucleotides, the RMSD value was found to be higher than 2 Å, indicating a large global conformational change (Fig. 2c and Extended Data Table 2).

To determine the cause of the global conformational change, we aligned the DBD and AdD domains of LIG1^EE/AA^ individually for 5’-dG:C and 5’-rG:C structures. The overlay of DBD demonstrates a conformational change in the AdD domain at the loop 1 (541-549 aa), and the overlay of AdD domain reveals a conformational change in the OBD domain at the loop 2 (749-759 aa) (Fig. 2e,f). These subtle changes observed in the loop 1 and loop 2 result in the large global conformational change because of the misalignment of entire domains relative to each other. A similar structural rearrangement has been previously reported for LIG4 undergoing a conformational change from an “open” LIG-AMP state to “closed” DNA-AMP conformation^49^.

Additionally, we aligned the AdD domain of LIG1^EE/AA^ for 5’-dG:C and 5’-rG:C structures and demonstrated that loop 3 (733-742 aa), which contains Lys(R)738 that shifts more than 10 Å closer to the 5’-P in the 5’-rG:C structure compared to the 5’-dG:C structure and forms a putative hydrogen bond with the 5’-P (Fig. 2g). We posit that this additional interaction deters AMP transfer from the ligase active site (K568) to the 5’-end of the nick (step 2) by increasing the energetic barrier to the adenylate transfer.

Lastly, the overlay of LIG1^EE/AA^/5’-rG:C (step 1) with all LIG1^WT^ or LIG1^EE/AA^ structures for 3’-RNA-DNA complexes and canonical DNA (steps 2 and step 3) as well as LIG1^EE/AA^ with a 3’-dA:C mismatch^36^ (PDB: 7SX5, step 1), revealed a difference in the DNA conformation (Fig. 3a). The RMSD using main chain atoms of LIG1^EE/AA^ for 5’-rG:C with 5’-dG:C and 3’-dA:C structures was found to be ~ 2.7Å (Extended Data Table 2). The main difference is the rotation of the 5’-P downstream of the nick DNA which widens the major groove downstream of the nick DNA and results in a narrowing of the minor groove (Fig. 3b,c). The minor groove is the interaction site of the OBD domain and narrowing the minor groove changes the conformation of OBD between the LIG1^EE/AA^/5’-dG:C and 5’-rG:C structures (Fig. 3d).

## Ligation efficiency of nick DNA containing a ribonucleotide at 3’-or 5’-end by LIG1

We finally tested the ligation efficiency of LIG1 using the nick DNA substrates with preinserted 3’-ribonucleotides and 5’-rG:C (Extended Data Scheme 2). Our results demonstrated the mutagenic ligation of the nick DNA substrates containing 3’-rA:T, 3’-rC:G, and 3’-rG:C (Fig. 4a). As expected, we obtained efficient nick sealing of canonical 3’-dA:T, 3’-dC:G, and 3’-dG:C ends (Extended Data Fig. 4) demonstrating no major difference in the amount of ligation products (Fig. 4b). On the other hand, our results demonstrated a lack of nick sealing by LIG1 in the presence of 5’-rG:C for the time points (10-60 sec) where we obtained efficient ligation of 3’-ribonucleotides (Fig. 4c, lanes 4-7) showing ~100-fold difference in the ligation products between 3’-ribonucleotides and 5’-rG:C (Fig. 4d). Consistent with previous report^47^, we observed ligase failure products with 5’-AMP-RNA for longer time points of the reaction (Fig. 4c, lanes 8-11 and Extended Data Fig. 5).

**Fig. 4.**
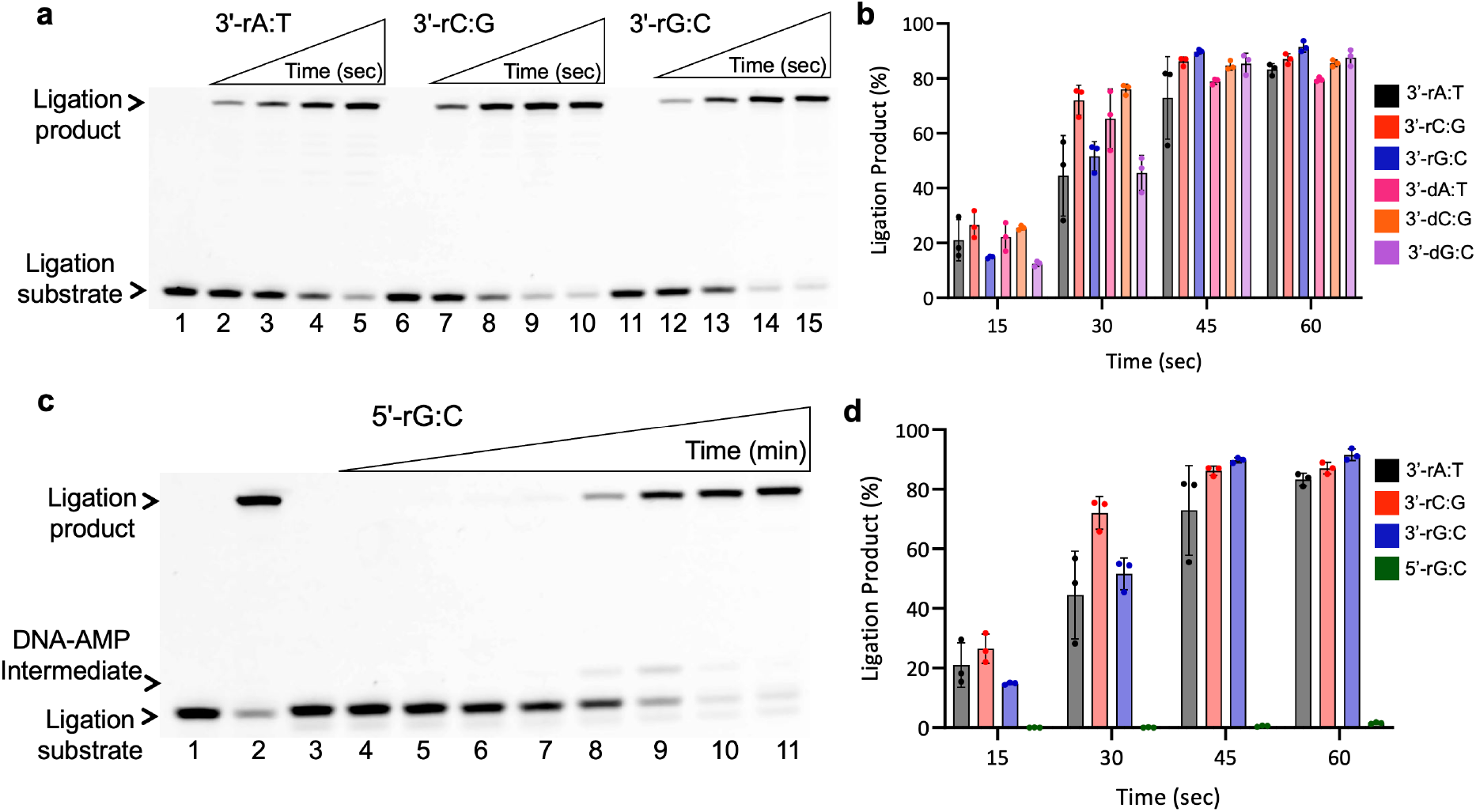
Ligation of the nick DNA with a ribonucleotide at 3’-or 5’-end by LIG1. **a,** Lanes 1, 6 and 11 are the negative enzyme controls of the nick DNA substrates with 3’-rA:T, 3’-rC:G, and 3’-rG:C, respectively. Lanes 2-5, 7-10, and 12-15 are the ligation products in the presence of 3’-rA:T, 3’-rC:G, and 3’-rG:C, respectively, and correspond to time points of 10, 30, 45, and 60 sec. **b,** Graph shows time-dependent change in the amount of ligation products for nick DNA substrates containing 3’-ribonucleotides *versus* 3’-deoxyribonucleotides. The data represent the average from three independent experiments ± SD. **c,** Line 1 is the negative enzyme control of the nick DNA substrate with 5’-dG:C, and line 2 is the positive control showing the ligation product in the presence of 5’-dG:C. Line 3 is the negative enzyme control of the nick DNA substrate with 5’-rG:C. Lanes 4-11 are the reaction products in the presence of 5’-rG:C, and correspond to time points of 0.25, 0.5, 0.75, 1, 5, 15, 30, and 60 min. **d,** Graph shows time-dependent change in the amount of ligation products for nick DNA substrates containing 3’-ribonucleotides *versus* 5’-rG:C. The data represent the average from three independent experiments ± SD.

## Discussion

LIG1 is responsible for the ligation of over 10 million Okazaki fragments during each round of DNA replication and nick sealing of the repair products at the final step of all DNA repair pathways^1–4^. Previously solved structures of LIG1/nick DNA duplexes with canonical, damaged, and mismatched ends have provided considerable details about steps 1 and 2 of the ligation reaction while relatively less information has been reported for final step^32–36^. Our structures uncover the features of 3’-RNA-DNA substrate and LIG1 interaction resulting in mutagenic outcomes at atomic resolution, which was entirely missing. This lesion mimics a DNA repair intermediate with 3’-ribonucleotides embedded in the genomic DNA because of their misincorporation by replicative and repair DNA polymerases in nucleus or mitochondria^11–27^. Using EDTA pre-treated LIG1^EE/AA^ protein, we captured the ligase active site while engaging with 3’-ribonucleotides (rA, rG, rC) during steps 2 and 3 in the crystal conditions lacking Mg^2+^. The lack of Mg^2+^ is confirmed in LIG1^WT^ structures for 3’-dA:T and 3’-rA:T, which contain the Mg^HiFi^ site, but show no density for Mg^2+^ (Extended Data Fig. 6). Previously, in the structure of ATP-dependent ligase from *Prochlorococcus marinus* (Pmar-Lig), the phosphodiester bond formation referring to step 3 of ligation reaction has been reported^50^. In B-form DNA, the sugar moieties usually adopt the 2’-endo sugar pucker conformation, while in A-form DNA or RNA duplexes the sugar moieties usually adopt the 3’-endo sugar pucker conformation^51^. The ligase active site forces the DNA upstream of the nick to adopt an A-form conformation which causes the sugar at the 3’-end to assume the 3’-endo conformation. Mg^2+^ can activate the 3’-OH for nucleophilic attack on the 5’-P and the 3’-end of nick DNA can also stabilize the 3’-endo conformation. We suggest that the LIG1^EE/AA^ structures for 3’-rG:C and 3’-rA:T that we captured in step 3 of the ligation reaction without Mg^2+^ in the crystal conditions is due to the 3’-endo conformation of sugar pucker that ribonucleotides naturally adopt.

Our study also contributes to the understanding of the mechanism by which LIG1 discriminates against a wrong sugar at the 5’-end of nick DNA. This 5’-RNA-DNA junction mimics the lesion could be inserted into DNA during RNase H2-dependent excision repair and trigger RNA-DNA damage response as previously reported^45–48^. In the LIG1/5’-RNA-DNA structure, we found a different closed state due to the conformational change of the DNA caused the ribonucleotide at the 5’-end. The OBD domain of LIG1 normally binds in the minor groove downstream of the nick DNA; however, in the LIG1/5’-rG:C structure, we observed a narrowing of the minor groove which initializes a major global conformational change in LIG1. Although it has been extensively reported that most DNA polymerases exhibit a “steric gate” (*i.e*., B-, and Y-family pols) or a protein backbone segment in minor groove nucleotide binding pocket (*i.e*, X-family pols) which governs ribonucleotide exclusion^16–27^, our study represents the first solved structure of RNA-DNA nick complexes for a human DNA ligase. Overall, our structures of LIG1 containing a single ribonucleotide at the 3’-or 5’-end of nick DNA contribute to understanding the molecular mechanism which governs ribonucleotide selectivity by a human ligase, LIG1 which functions in nuclear replication and repair. Future studies with other human DNA ligases will be aimed at untangling the impact of improper sugar on the ligation reaction to gain a deeper understanding of how fidelity of nick sealing is ensured to maintain genome integrity.

## Methods

### Protein purifications

Recombinant proteins with 6x-his tag for the C-terminal (Δ261) wild-type and E346A/E592A (EE/AA) mutant of human DNA ligase I (LIG1) were overexpressed in Rosetta (DE3) *E. coli* cells in Terrific Broth (TB) media with kanamycin (50 μgml^-1^) and chloramphenicol (34 μgml^-1^) at 37 °C^52–58^. The cells were induced with 0.5 mM isopropyl β-D-thiogalactoside (IPTG) when the OD_600_ was reached to 1.0, and the overexpression was continued for overnight at 28 °C. Cells were lysed in the lysis buffer containing 50 mM Tris-HCl (pH 7.0), 500 mM NaCl, 20 mM imidazole, 10% glycerol, 1 mM PMSF, an EDTA-free protease inhibitor cocktail tablet by sonication at 4 °C. The cell lysate was pelleted at 31,000 x g for 1 h at 4 °C. Proteins were purified by a HisTrap HP column with an increasing imidazole concentration (20-300 mM) after being equilibrated in the binding buffer containing 50 mM Tris-HCl (pH 7.0), 500 mM NaCl, 20 mM imidazole, 10% glycerol at 4 °C. The collected fractions were subsequently loaded onto a HiTrap Heparin column that was equilibrated with the binding buffer containing 50 mM Tris-HCl (pH 7.0), 50 mM NaCl, 1.0 mM EDTA, and 10% glycerol, and then eluted with a linear gradient of NaCl up to 1 M. LIG1 proteins were further purified by Superdex 200 10/300 column in the buffer containing 20 mM Tris-HCl (pH 7.0), 200 mM NaCl, and 1 mM DTT. All proteins concentrated, frozen in liquid nitrogen, and stored at −80 °C.

### Crystallization and structure determination

Nick DNA substrates for LIG1 X-ray crystallography were prepared by annealing upstream, downstream, and template primers (Extended Data Table 4). LIG1 C-terminal (Δ261) wild-type (LIG1^WT^) and EE/AA mutant (LIG1^EE/AA^) proteins were pre-treated with EDTA before using in the crystallization experiments in the absence of Mg^2+^ as we reported previously^36^. All LIG1-nick DNA complex crystals were grown at 20 °C using the hanging drop method. LIG1 (at 27 mgml^-1^)/DNA complex solution was prepared in 20 mM Tris-HCl (pH 7.0), 200 mM NaCl, 1 mM DTT, 1 mM EDTA and 1 mM ATP at 1:4:1 DNA:protein molar ratio and then mixed with 1 μl reservoir solution. The crystal conditions are described in Extended Data Table 5. Crystals were harvested and submerged in cryoprotectant solution containing reservoir solution mixed with glycerol to a final concentration of 20% glycerol before being flash cooled in liquid nitrogen for data collection (HKL Research, Inc). Crystals were kept at 100 °K during X-ray diffraction data collection using the beamlines APS-22-ID and CHESS-7B2. All structures were solved by the molecular replacement method using PHASER with PDB entry 7SUM as a search model^59^. Iterative rounds of model building were performed in COOT and the final models were refined with PHENIX or REFMAC5^60–62^. 3DNA program was used for sugar pucker analysis^63^. All structural images were drawn using PyMOL (The PyMOL Molecular Graphics System, V0.99, Schrödinger, LLC). Detailed crystallographic statistics are provided in Extended Data Table 1.

### DNA ligation assays

Nick DNA substrates with a preinserted ribonucleotide at 3’-or 5’-end were prepared to evaluate the ligation efficiency of LIG1 in the presence of 3’-rA:T, 3’-rC:G, 3’-rG:C or 5’-rG:C (Extended Data Scheme 2). In addition, we tested the nick DNA substrates with cognate ends 3’-dA:T, 3’-dC:G, 3’-dG:C and 5’-dG:C (Extended Data Table 6). The reaction containing 50 mM Tris-HCl (pH: 7.5), 100 mM KCl, 10 mM MgCl2, 1 mM ATP, 1 mM DTT, 100 μgml^-1^ BSA, 10% glycerol, and nick DNA substrate (500 nM) was initiated by the addition of LIG1 (100 nM). The reaction samples were incubated at 37 °C, stopped by quenching with an equal amount of the buffer containing 95% formamide, 20 mM ethylenediaminetetraacetic acid, 0.02% bromophenol blue and 0.02% xylene cyanol, and collected at the time points indicated in the figure legends. The reaction products were then separated by electrophoresis on an 18% denaturing polyacrylamide gel. The gels were scanned with a Typhoon Phosphor Imager (Amersham Typhoon RGB), and the data were analyzed using ImageQuant software^52–58^.

## Data availability

Atomic coordinates and structure factors for the reported crystal structure of LIG1 in complex with nick DNA complexes has been deposited in the RCSB Protein Data Bank under accession numbers 8FYK, 8FZX, 8FZL, 8G1X, 8FZS, 8G0J, 8G1O, 8G11, 8G37. All relevant data are available from the authors upon reasonable request.

## Acknowledgements

This work is based upon research conducted at the Center for High Energy X-ray Sciences (CHEXS), which is supported by the National Science Foundation under award DMR-1829070, and the Macromolecular Diffraction at CHESS (MacCHESS) facility, which is supported by award 1-P30-GM124166-01A1 from the National Institute of General Medical Sciences, National Institutes of Health, and by New York State’s Empire State Development Corporation (NYSTAR). This research used resources of the Advanced Photon Source (APS), U.S. Department of Energy (DOE) Office of Science user facility operated for the DOE Office of Science by Argonne National Laboratory under Contract No. DE-AC02-06CH11357. The authors thank to Dr. Craig Vander Kooi (University of Florida) for his exceptional support with the APS beam time and data collection.

## Author contributions

Conceptualization M.Ç., methodology and investigation T.Q., M.G., M.Ç.; writing-original draft T.Q., M.G., M.Ç.; writing-reviewing and editing T.Q., M.G., M.Ç.; funding acquisition M.Ç.

