## supplementary information for "Structures of LIG1 uncover the mechanism of sugar discrimination against a ribonucleotide at 3’- and 5’-end of the nick DNA"

Extended Data Figures 1-6

Extended Data Tables 1-6

Extended Data Schemes 1-2

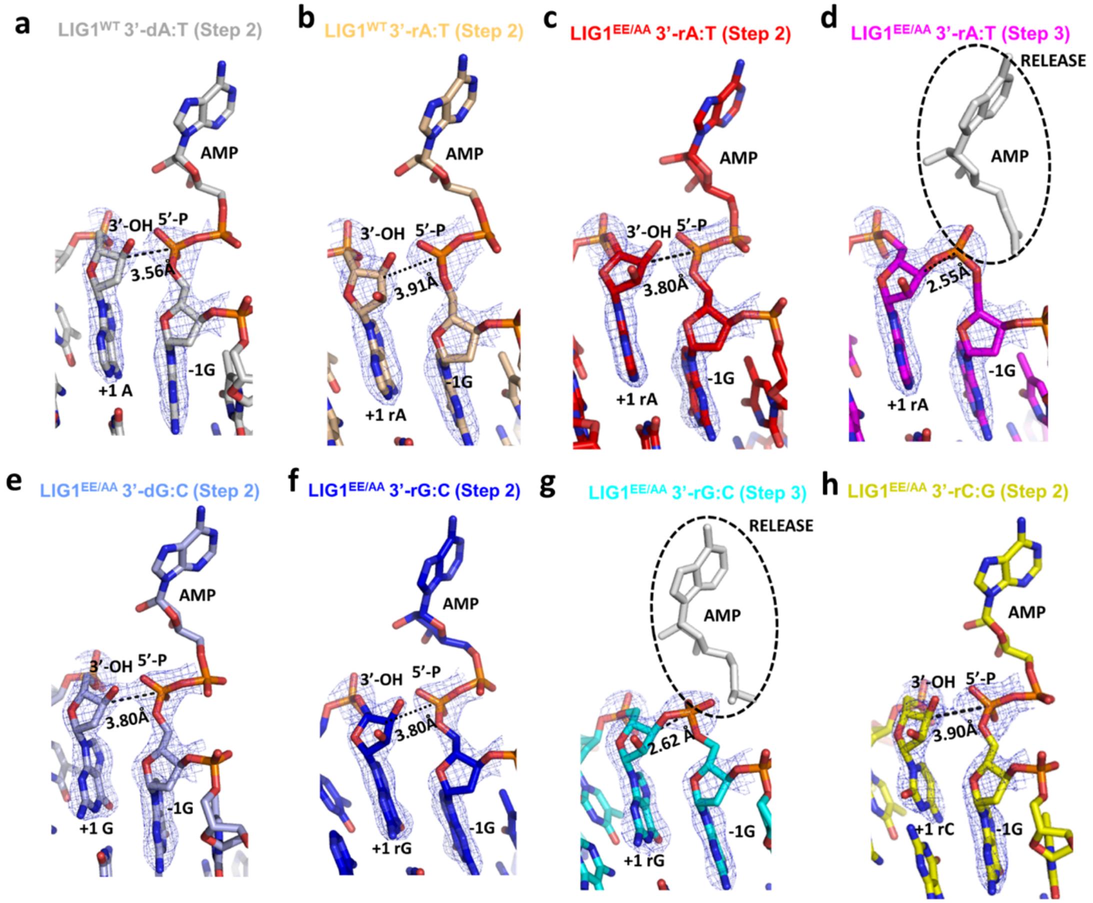


**Extended Data Fig. 1. LIG1 structures shows the differences in the distances between 3'- and 5'- terminus of the nick DNA.** Structures of LIG1$^{WT}$ and LIG1$^{EE/AA}$ in complex with nick DNA containing ribonucleotide or deoxyribonucleotide at the 3'-end of upstream show for 3'-dA:T (**a**, step 2 for wild-type), 3'-rA:T (**b**, step 2 for wild-type), 3'-rA:T (**c**, step 2 for EE/AA), 3'-rA:T (**d**, step 3 for EE/AA), 3'-dG:C (**e**, step 2 for wild-type), 3'-rG:C (**f**, step 2 for EE/AA), 3'-rG:C (**g**, step 3 for EE/AA), 3'-rC:G (**h**, step 2 for EE/AA). 2Fo - Fc density map of 3'- and 5'- terminus of the nick DNA substrates are contoured at 3σ. In the LIG1 structures solved in the final step 3 of the ligation reaction (d, g), the maps of AMPs are incomplete for AMP. The distances are shown between C3' atom of the sugar on the 3'- and 5'-P ends of the nick DNA.

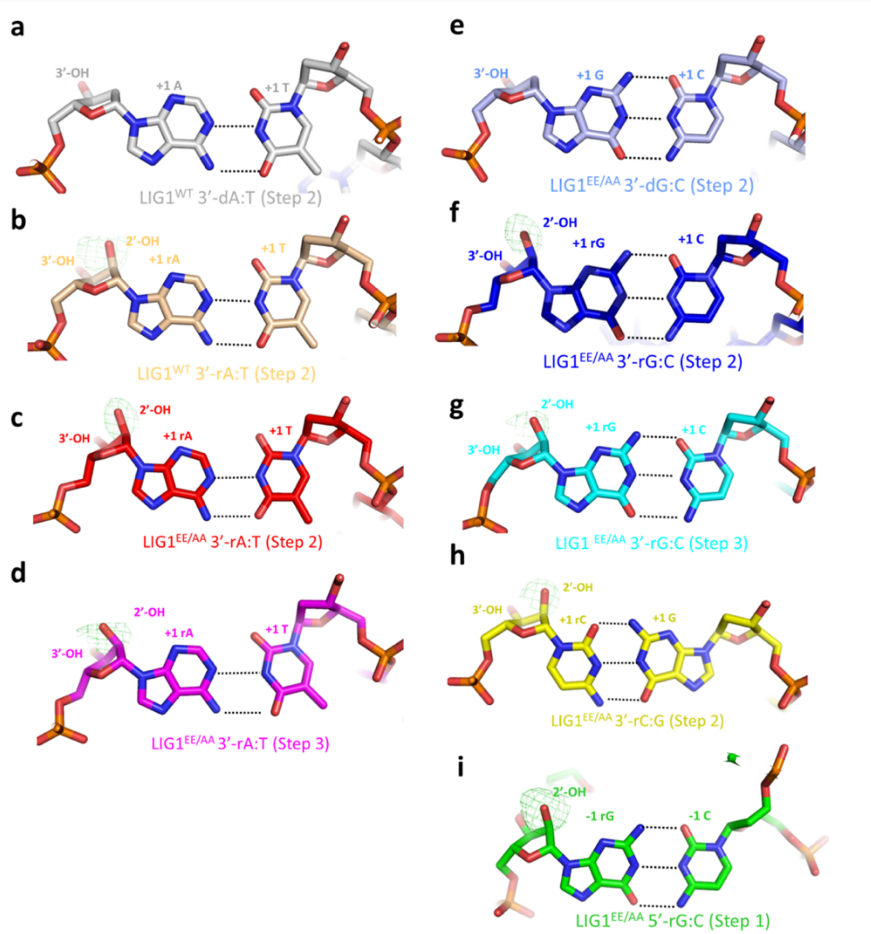


**Extended Data Fig. 2. Base-pairing architecture of ribonucleotide-containing DNA ends at the 3'-OH terminus.** Watson-crick base pairing between 3'- and 5'- terminus of the nick DNA are shown for LIG1$^{WT}$/3'-dA:T (**a**), LIG1$^{WT}$/3'-rA:T (**b**), LIG1$^{EE/AA}$/3'-rA:T (**c**, step 2), LIG1$^{EE/AA}$/3'-rA:T (**d**, step 3), LIG1$^{EE/AA}$/3'-dG:C (**e**), LIG1$^{EE/AA}$/3'-rG:C (**f**), LIG1$^{EE/AA}$/3'-rG:C (**g**), LIG1$^{EE/AA}$/3'-rC:G (**h**) and LIG1$^{EE/AA}$/5'-rG:C (**i**). Simulated annealing omit maps (Fo-Fc) of the 2'-OH atom of ribose are contoured at 3σ.

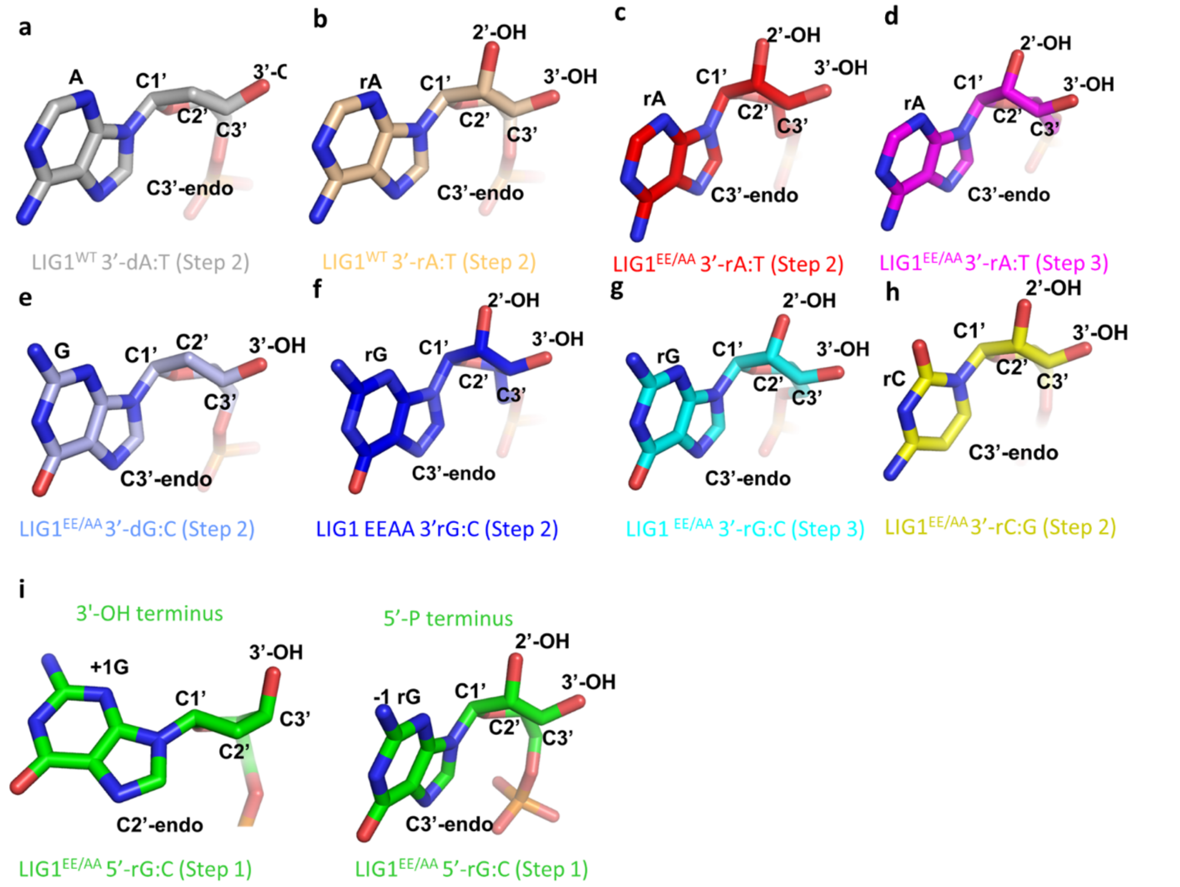


**Extended Data Fig. 3. Sugar pucker analyses of LIG1 structures solved in the present study.** 3'-end sugar pucker analyses of LIG1 structures are presented for LIG1$^{WT}$/3'-dA:T (**a**), LIG1$^{WT}$/3'-rA:T (**b**), LIG1$^{EE/AA}$/3'-rA:T (**c**, step 2), LIG1$^{EE/AA}$/3'-rA:T (**d**, step 3), LIG1$^{EE/AA}$/3'-dG:C (**e**), LIG1$^{EE/AA}$/3'-rG:C (**f**, step 2), LIG1$^{EE/AA}$/3'-rG:C (**g**, step 3), LIG1$^{EE/AA}$/3'-rC:G (**h**) LIG1$^{EE/AA}$/5'-rG:C (**i**), and for 5'-P end of LIG1$^{EE/AA}$/5'-rG:C.

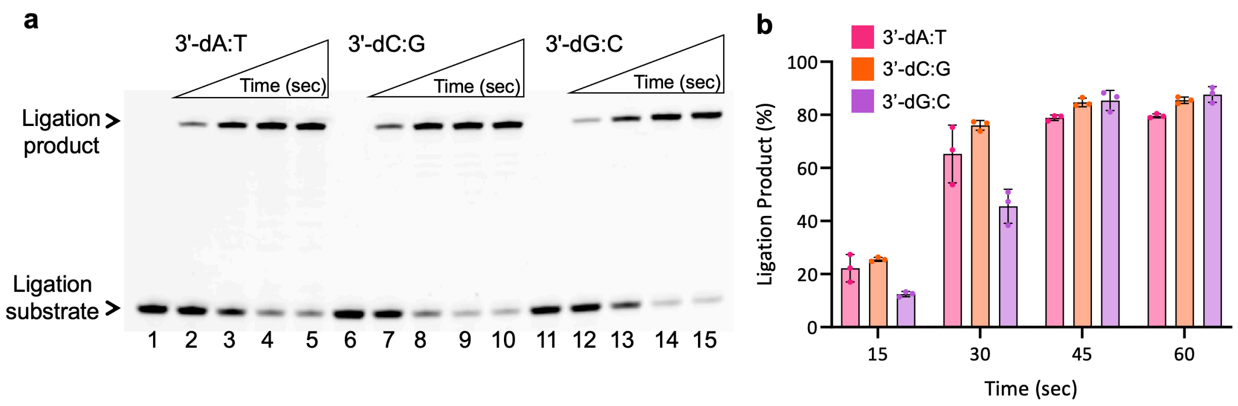


**Extended Data Fig. 4. Ligation of the nick DNA with cognate base pairs by LIG1. a,** Lanes 1, 6 and 11 are the negative enzyme controls of the nick DNA substrates with 3'-dA:T, 3'-dC:G, and 3'-dG:C, respectively. Lanes 2-5, 7-10, and 12-15 are the ligation products in the presence of 3'-dA:T, 3'-dC:G, and 3'-dG:C, respectively, and correspond to time points of 10, 30, 45, and 60 sec. **b,** Graph shows time-dependent change in the amount of ligation products for nick DNA substrates containing 3'-deoxyribonucleotides. The data represent the average from three independent experiments ± SD.

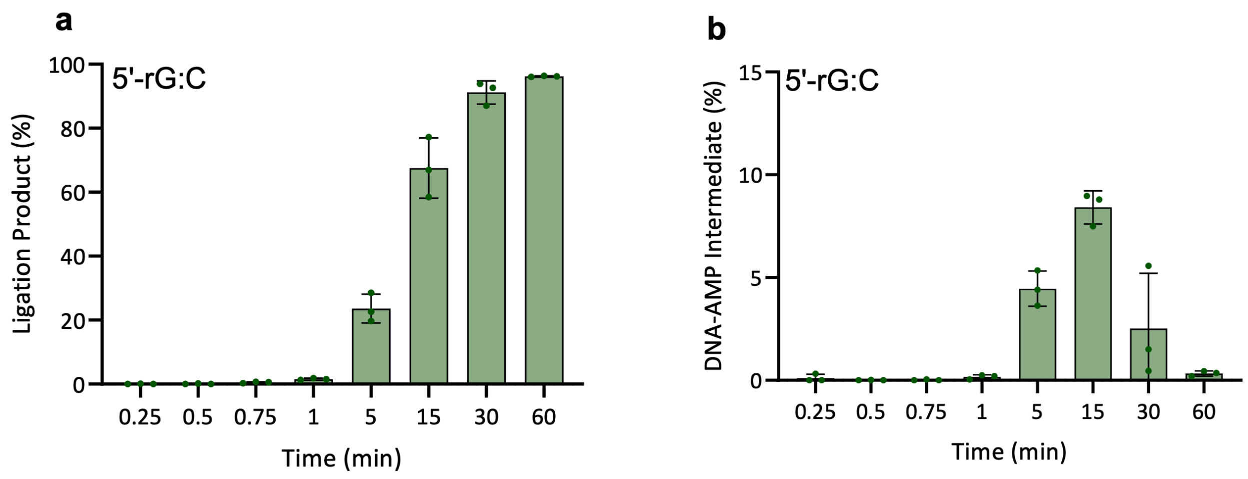


**Extended Data Fig. 5. Ligation of the nick DNA with 5'-ribonucleotide by LIG1.** Graphs show time-dependent change in the amount of ligation (**a**) and ligation failure or DNA-AMP intermediate (**b**) products for the nick DNA substrate containing 5'-rG:C. The data represent the average from three independent experiments ± SD.

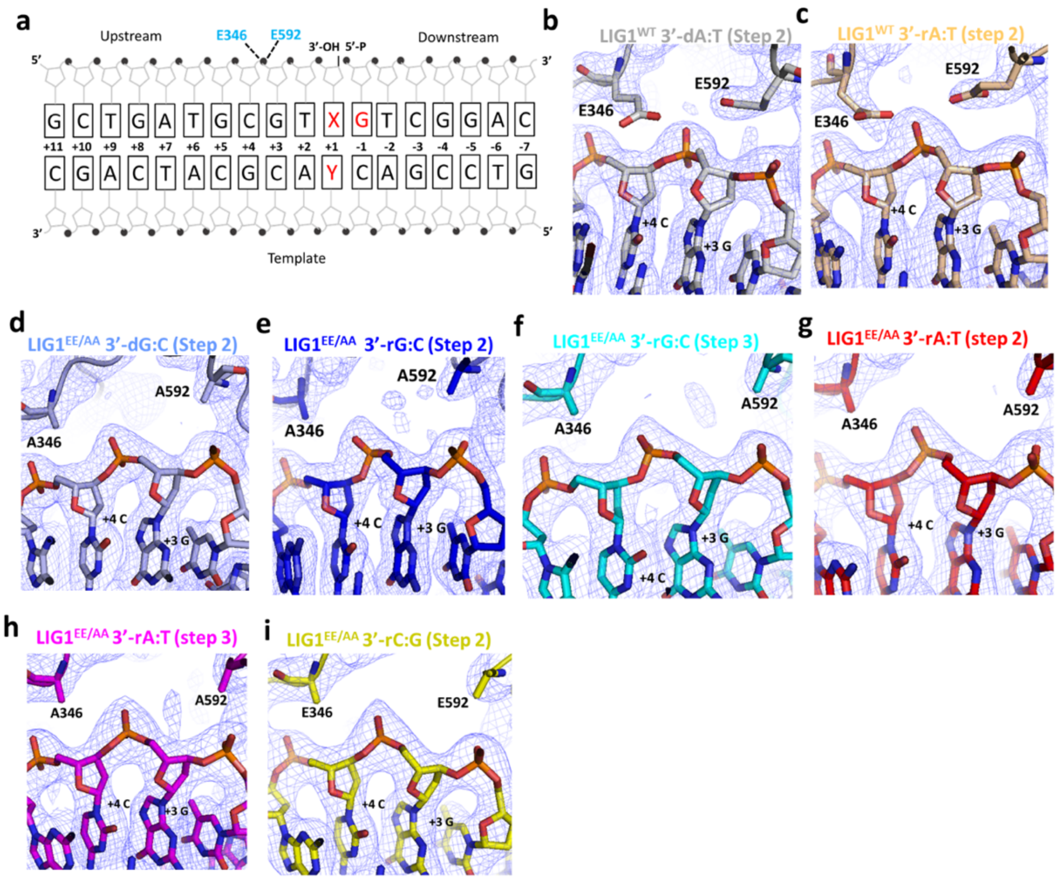


**Extended Data Fig. 6. Structures of LIG1$^{WT}$ and LIG1$^{EE/AA}$ demonstrate the position of high-fidelity site. a**, Schematic view of nick DNA showing the amino acids E346 and E592 that are identified as a high-fidelity and $Mg^{2+}$ binding site in the previously solved LIG1 structures. **b-i**, The mutations at this side (E346/A346 and E592/A592) are shown in the structures of LIG1$^{WT}$/3'-dA:T (step 2, **b**), LIG1$^{WT}$/3'-rA:T (step 2, **c**), LIG1$^{EE/AA}$/3'-dG:C (step 2, **d**) LIG1$^{EE/AA}$/3'-rG:C (step 2, **e**, step 2), LIG1$^{EE/AA}$/3'-rG:C (**f**, step 3), LIG1$^{EE/AA}$/3'-rA:T (step 2, **g**), LIG1$^{EE/AA}$/3'-rA:T (step 3, **h**), LIG1$^{EE/AA}$/3'-rC:G (step 2, **i**). 2Fo - Fc density maps are contoured at 1σ. E346/A346, E592/A592 and DNA are shown as sticks.

**a**

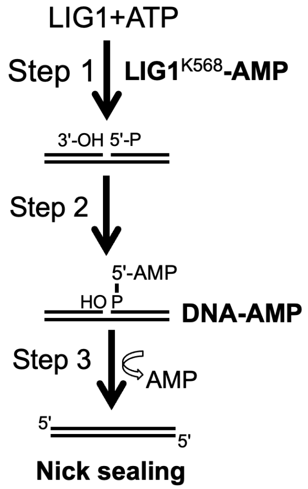


**b**

| Protein | LIG1 structure | Ligation step | Reference |
|---|---|---|---|
| $LIG1^{WT}$ | 3'-ddC:G | Step 2 | 32 |
| $LIG1^{WT}$ | 3'-ddC:G+$Mg^{2+}$ | Step 2 | 33 |
| $LIG1^{WT}$ | 3'-dC:G | Step 2 | 33 |
| $LIG1^{EE/AA}$ | 3'-dC:G | Step 2 | 33 |
| $LIG1^{EE/AA}$ | 3'-8oxodG:A | Step 2 | 33 |
| $LIG1^{EE/AA}$ | Bulged DNA | Step 2 | 34 |
| $LIG1^{R641L}$ | 3'-dC:G | Step 2 | 35 |
| $LIG1^{R771W}$ | 3'-dC:G | Step 2 | 35 |
| $LIG1^{EE/AA}$ | 3'-dA:T | Step 2 | 36 |
| $LIG1^{EE/AA}$ | 3'-dG:T | Step 2 | 36 |
| $LIG1^{EE/AA}$ | 3'-dA:C | Step 1 | 36 |
| $LIG1^{EE/AA}$ | 5'-rG:C | Step 1 | Present study |
| $LIG1^{WT}$ | 3'-dA:T | Step 2 | Present study |
| $LIG1^{EE/AA}$ | 3'-dG:C | Step 2 | Present study |
| $LIG1^{WT}$ | 3'-rA:T | Step 2 | Present study |
| $LIG1^{EE/AA}$ | 3'-rA:T | Step 2 | Present study |
| $LIG1^{EE/AA}$ | 3'-rC:G | Step 2 | Present study |
| $LIG1^{EE/AA}$ | 3'-rG:C | Step 2 | Present study |
| $LIG1^{EE/AA}$ | 3'-rA:T | Step 3 | Present study |
| $LIG1^{EE/AA}$ | 3'-rG:C | Step 3 | Present study |

**Extended Data Scheme 1. Illustration of DNA ligation reaction. a,** Three steps of ligation reaction involves the initial formation of DNA ligase-AMP intermediate when K568 active site of LIG1 is adenylated in step 1, the transfer of AMP to 5'-P end of nick DNA resulting in the formation of DNA-AMP intermediate in step 2, and final phosphodiester bond formation coupled to AMP release in step 3. **b**, List of LIG1 structures in complex with nick DNA complexes with canonical, damaged, mismatched, or ribonucleotide containing ends.

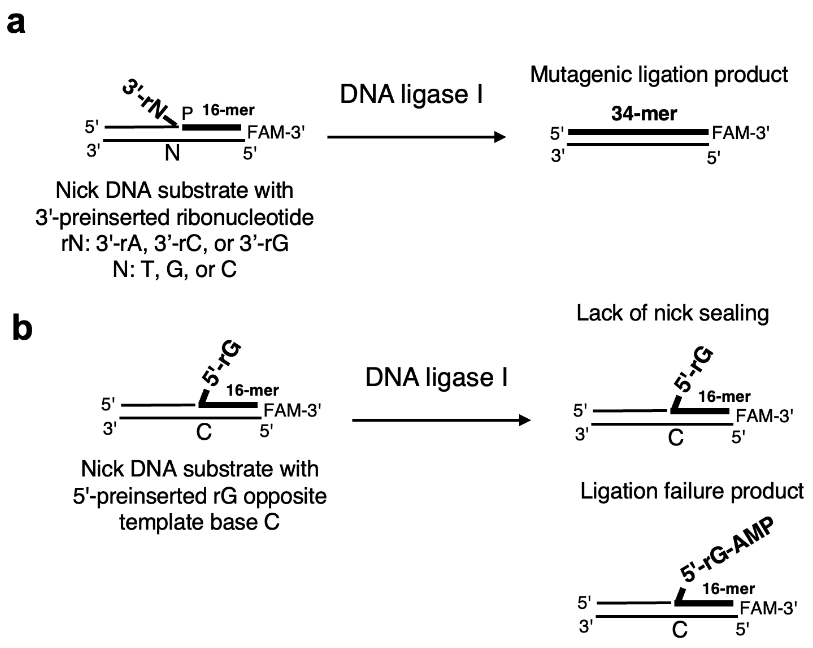


**Extended Data Scheme 2. Illustrations of nick sealing assays used in this study**. Ligation assays were used to investigate the sugar discrimination mechanism of LIG1 for the nick DNA substrates with preinserted 3'-ribonucleotides 3'-rA:T, 3'-rC:G, 3'-rG:C (**a**) and 5'-rG:C (**b**).

**Supplementary Table 1.** X-ray data collection and refinement statistics

| | LIG1$^{WT}$ 3'-dA:T | LIG1$^{WT}$ 3'-rA:T | LIG1$^{EE/AA}$ 3'-rA:T (Step 2) | LIG1$^{EE/AA}$ 3'-rA:T (Step 3) | LIG1$^{EE/AA}$ 3'-dG:C | LIG1$^{EE/AA}$ 3'-rG:C (Step 2) | LIG1$^{EE/AA}$ 3'-rG:C (Step 3) | LIG1$^{EE/AA}$ 3'-rC:G | LIG1$^{EE/AA}$ 5'-rG:C |
|---|---|---|---|---|---|---|---|---|---|
| PDB entry ID | **8FYK** | **8FZX** | **8FZL** | **8G1X** | **8FZS** | **8G0J** | **8G1O** | **8G11** | **8G37** |
| **Data collection** | | | | | | | | | |
| Space group | $P2_12_12_1$ | $P2_12_12_1$ | $P2_12_12_1$ | $P2_12_12_1$ | $P2_12_12_1$ | $P2_12_12_1$ | $P2_12_12_1$ | $P2_12_12_1$ | $P2_1$ |
| Cell | | | | | | | | | |
| *a*, *b*, *c* (Å) | 64.4 115.4 125.9 | 64.5 115.6 126.0 | 64.8 115.9 123.0 | 64.1 115.3 125.0 | 65.0 115.6 126.1 | 65.1 115.3 123.9 | 64.7 115.9 126.2 | 64.3 115.6 125.5 | 71.3 117.2 101.5 |
| α, β, γ (°) | 90 | 90 | 90 | 90 | 90 | 90 | 90 | 90 | 90.0 96.7 90.0 |
| Resolution (Å) | 20-2.8 (2.85-2.8) | 20-2.95 (3.0-2.95) | 20-2.54 (2.58-2.54) | 20-2.9 (2.95-2.9) | 20-2.51 (2.55-2.51) | 20-2.45 (2.49-2.45) | 20-2.8 (2.85-2.8) | 20-2.69 (2.74-2.69) | 20-2.4 (2.44-2.4) |
| $R_{sym}$ | 0.136 (0.920) | 0.078 (1.089) | 0.112 (0.791) | 0.075 (0.544) | 0.068 (0.779) | 0.117 (0.850) | 0.076 (1.032) | 0.123 (0.671) | 0.159 (0.998) |
| $I / \sigma (I)$ | 19.4 (1.9) | 18.1 (1.2) | 27.7 (1.8) | 25.8 (2.2) | 37.2 (1.7) | 28.6 (1.7) | 44.0 (2.9) | 31.2 (1.8) | 17.0 (1.6) |
| $CC_{1/2}$ | 0.957 (0.791) | 0.992 (0.702) | 0.988 (0.726) | 0.994 (0.882) | 0.996 (0.803) | 0.979 (0.651) | 0.992 (0.863) | 0.973 (0.823) | 0.981 (0.599) |
| $CC^*$ | 0.989 (0.940) | 0.998 (0.908) | 0.997 (0.917) | 0.998 (0.968) | 0.999 (0.944) | 0.995 (0.888) | 0.998 (0.963) | 0.993 (0.950) | 0.995 (0.865) |
| Completeness | 99.7 (99.7) | 99.1 (99.9) | 99.7 (100.0) | 98.4 (99.6) | 99.8 (99.9) | 98.8 (95.9) | 99.8 (100.0) | 99.7 (99.9) | 99.9 (99.8) |
| Redundancy | 6.5 (6.5) | 6.5 (7.1) | 6.2 (6.2) | 6.0 (6.8) | 6.5 (6.7) | 6.6 (5.8) | 12.4 (13.6) | 6.5 (5.6) | 6.7 (6.1) |
| **Refinement** | | | | | | | | | |
| Resolution (Å) | 20-2.80 | 20-2.95 | 20-2.54 | 20-2.9 | 20-2.51 | 20-2.45 | 20-2.80 | 20-2.69 | 20-2.4 |
| No. reflections | 23408 | 20076 | 30575 | 20558 | 31507 | 34363 | 22466 | 26130 | 61275 |
| $R_{work}/R_{free}$ | 18.6/21.6 | 19.6/24.8 | 17.5/19.7 | 17.4/22.2 | 17.7/20.2 | 17.4/20.9 | 19.0/24.4 | 17.6/20.9 | 18.4/23.8 |
| Non-H atoms: | 5738 | 5663 | 5824 | 5590 | 5733 | 5876 | 5721 | 5647 | 10636 |
| Protein | 4868 | 4899 | 4957 | 4824 | 4937 | 4973 | 4901 | 4871 | 8630 |
| DNA/RNA | 733 | 734 | 734 | 733 | 733 | 734 | 733 | 734 | 1474 |
| AMP | 22 | 22 | 22 | | 22 | 22 | | 22 | 44 |
| $H_2O$ | 115 | 8 | 111 | 33 | 41 | 147 | 87 | 20 | 480 |
| Average B- | 69.7 | 81.1 | 62.8 | 81.4 | 78.5 | 57.2 | 40.9 | 77.9 | 41.8 |
| Protein | 73.1 | 84.5 | 66.0 | 84.4 | 82.5 | 59.7 | 43.8 | 81.5 | 41.3 |
| DNA | 49.8 | 58.5 | 44.2 | 54.6 | 53.8 | 42.3 | 23.3 | 53.8 | 44.5 |

| | | | | | | | | | |
|---|---|---|---|---|---|---|---|---|---|
| AMP | 74.5 | 86.5 | 56.0 | | 43.4 | 48.7 | | 80.4 | 31.0 |
| $H_2O$ | 49.4 | 61.3 | 41.5 | 66.7 | 70.00 | 48.2 | 32.1 | 64.7 | 43.1 |
| R.M.S.D | | | | | | | | | |
| Bond lengths | 0.011 | 0.012 | 0.011 | 0.011 | 0.010 | 0.009 | 0.011 | 0.009 | 0.010 |
| Bond angles (°) | 1.204 | 1.405 | 1.198 | 1.283 | 1.179 | 1.076 | 1.288 | 1.135 | 1.145 |

| **R.M.S.D (Å)** | LIG1$^{WT}$ 3'-dA:T | LIG1$^{WT}$ 3'-rA:T | LIG1$^{EE/AA}$ 3'-rA:T (Step 2) | LIG1$^{EE/AA}$ 3'-rA:T (Step 3) | LIG1$^{EE/AA}$ 3'-dG:C | LIG1$^{EE/AA}$ 3'-rG:C (Step 2) | LIG1$^{EE/AA}$ 3'-rG:C (Step 3) | LIG1$^{EE/AA}$ 3'-rC:G | LIG1$^{EE/AA}$ 5'-rG:C |
|---|---|---|---|---|---|---|---|---|---|
| LIG1$^{WT}$ 3'-dA:T | | 0.381 | 0.565 | 0.463 | 0.369 | 0.545 | 0.371 | 0.364 | 2.061 |
| LIG1$^{WT}$ 3'-rA:T | | | 0.550 | 0.396 | 0.347 | 0.510 | 0.303 | 0.284 | 2.041 |
| LIG1$^{EEAA}$ 3'-rA:T (Step 2) | | | | 0.620 | 0.502 | 0.515 | 0.535 | 0.515 | 2.153 |
| LIG1$^{EEAA}$ 3'-rA:T (Step 3) | | | | | 0.439 | 0.595 | 0.393 | 0.382 | 2.028 |
| LIG1$^{EEAA}$ 3'-dG:C | | | | | | 0.454 | 0.327 | 0.309 | 2.037 |
| LIG1$^{EEAA}$ 3'-rG:C (Step 2) | | | | | | | 0.493 | 0.476 | 2.121 |
| LIG1$^{EEAA}$ 3'-rG:C (Step 3) | | | | | | | | 0.277 | 2.045 |
| LIG1$^{EEAA}$ 3'-rC:G | | | | | | | | | 2.048 |

**Extended Data Table 2.** The root mean square deviation (RMSD) of all LIG1 structures solved in the present study.

| R.M.S.D (Å) | DBD (262-535 aa) | AdD (536-748 aa) | AdD without loop1 and loop2 (550-730 aa) | OBD (749-900 aa) | OBD without loop 3 (760-900 aa) | DNA |
|---|---|---|---|---|---|---|
| LIG1$^{EE/AA}$ 5'-rG:C<br>LIG1$^{EE/AA}$ 5'-dG:C | 0.672 | 1.121 | 0.587 | 0.917 | 0.685 | 2.799 |
| LIG1$^{EE/AA}$ 5'-rG:C<br>LIG1$^{EE/AA}$ 3'-dA:C (7SUM) | 0.641 | 1.079 | 0.467 | 1.062 | 0.846 | 2.677 |

**Extended Data Table 3.** RMSD of LIG1$^{EE/AA}$/5'-rG:C with LIG1$^{EE/AA}$ 5'-dG/C or 3'-dA:C structures compared by all the ligase domains and nick DNA. DBD, AdD, and OBD are DNA-binding domain, Adenylation domain, and Oligonucleotide-binding domain, respectively.

| Oligonucleotide | Sequence (5'-3') |
|---|---|
| Template T | GTCCGAC<u>T</u>ACGCATCAGC |
| Template C | GTCCGAC<u>C</u>ACGCATCAGC |
| Template G | GTCCGAC<u>G</u>ACGCATCAGC |
| Upstream A (3'-rA) | GCTGATGCGT**A** |
| Upstream C (3'-rC) | GCTGATGCGT**C** |
| Upstream G (3'-rG) | GCTGATGCGT**G** |
| Downstream (5'-P) | P-GTCGGAC |
| Upstream G (3'-G) | GCTGATGCGTG |
| Downstream (5'-rG) | P-**G**TCGGAC |

**Extended Data Table 4. Oligonucleotides used for LIG1 crystallization.** Upstream oligonucleotides A, C, G including a single ribonucleotide at the 3'-end, downstream oligo with phosphate (P) at the 5'-end, and template oligonucleotides containing T, C, or G on a template position were used to prepare the nick DNA substrates with 3'-ribonucleotides (3'-dA:T, 3'-dC:G, and 3'-dG:C) for LIG1$^{WT}$ and LIG1$^{EE/AA}$ crystallizations. Upstream oligonucleotide G including a single deoxyribonucleotide at the 3'-end, downstream oligo with a single ribonucleotide at the 5'-end, and template oligonucleotide containing C on a template position were used to prepare the nick DNA substrate with 5'-ribonucleotide (5'-rG:C) for LIG1$^{EE/AA}$ crystallization. The base at template base position is underlined and the ribonucleotide base position is shown in bold.

| Protein/DNA | Crystal conditions | Crystalized time (day) |
|---|---|---|
| LIG1$^{WT}$ 3'-dA:T | 100 mM MES (pH 5.7), 100 mM lithium acetate, and 8% (w/v) PEG3350 | 1 |
| LIG1$^{WT}$ 3'-rA:T | 100 mM MES (pH 6.4), 100 mM lithium acetate, and 16% (w/v) PEG3350 | 1 |
| LIG1$^{EE/AA}$ 3'-rA:T (Step 2) | 100 mM MES (pH 6.7), 100 mM lithium acetate, and 16% (w/v) PEG3350 | 1 |
| LIG1$^{EE/AA}$ 3'-rA:T (Step 3) | 100 mM MES (pH 6.7), 100 mM lithium acetate, and 16% (w/v) PEG3350 | 3 |
| LIG1$^{EE/AA}$ 3'-dG:C | 100 mM MES (pH 6.1), 100 mM lithium acetate, and 10% (w/v) PEG3350 | 1 |
| LIG1$^{EE/AA}$ 3'-rG:C (Step 2) | 100 mM MES (pH 7.0), 100 mM lithium acetate, and 14% (w/v) PEG3350 | 1 |
| LIG1$^{EE/AA}$ 3'-rG:C (Step 3) | 100 mM MES (pH 7.0), 100 mM lithium acetate, and 14% (w/v) PEG3350 | 3 |
| LIG1$^{EE/AA}$ 3'-rC:G | 100 mM MES (pH 6.1), 250 mM lithium acetate, and 14% (w/v) PEG3350 | 1 |
| LIG1$^{EE/AA}$ 5'-rG:C | 100 mM MES (pH 6.8), 100 mM lithium acetate, and 20% (w/v) PEG3350 | 4 |

**Extended Data Table 5.** Crystalization conditions of LIG1$^{WT}$ and LIG1$^{EE/AA}$ structures solved in the present study.

| DNA Substrates | Sequence |
|---|---|
| Nick DNA with 3'-dA:T | 5'-CATGGGCGGCATGAACC**A**GAGGCCCATCCTCACC-3-FAM<br>3'-GTACCCGCCGTACTTGGTCTCCGGGTAGGAGTGG-5' |
| Nick DNA with 3'-dC:G | 5'-CATGGGCGGCATGAACC**C**GAGGCCCATCCTCACC-3-FAM<br>3'-GTACCCGCCGTACTTGGGCTCCGGGTAGGAGTGG-5' |
| Nick DNA with 3'-dG:C | 5'-CATGGGCGGCATGAACC**G**GAGGCCCATCCTCACC-3-FAM<br>3'-GTACCCGCCGTACTTGGCCTCCGGGTAGGAGTGG-5' |
| Nick DNA with 3'-rA:T | 5'-CATGGGCGGCATGAACC**A**GAGGCCCATCCTCACC-3-FAM<br>3'-GTACCCGCCGTACTTGGTCTCCGGGTAGGAGTGG-5' |
| Nick DNA with 3'-rC:G | 5'-CATGGGCGGCATGAACC**C**GAGGCCCATCCTCACC-3-FAM<br>3'-GTACCCGCCGTACTTGGGCTCCGGGTAGGAGTGG-5' |
| Nick DNA with 3'-rG:C | 5'-CATGGGCGGCATGAACC**C**GAGGCCCATCCTCACC-3-FAM<br>3'-GTACCCGCCGTACTTGGGCTCCGGGTAGGAGTGG-5' |
| Nick DNA with 5'-rG:C | 5'-CATGGGCGGCATGAACCG**G**AGGCCCATCCTCACC-3-FAM<br>3'-GTACCCGCCGTACTTGGCCTCCGGGTAGGAGTGG-5' |

**Extended Data Table 6. Nick DNA substrates used in ligation assays.** FAM denotes a fluorescence tag and is located at the 3'-end of the nick DNA substrates. The base at 3'-end is shown as bold and a template base is underlined.
